# The neural processes of illusory occlusion in object recognition

**DOI:** 10.1101/2025.05.28.656741

**Authors:** Almudena Ramírez-Haro, Denise Moerel, Genevieve L. Quek, Manuel Varlet, Tijl Grootswagers

## Abstract

Fast and accurate object recognition is crucial for effective behaviour in dynamic visual environments. In some cases, the visual system must overcome ambiguity in visual input during object recognition, such as when an object is partially hidden behind another. Recurrent processing between higher- and lower-order areas is thought to play a role in resolving such ambiguity, enabling the filling in of missing visual information. Here we examined this claim using a novel paradigm in which partial object images appear ‘occluded’ by an illusory Kanizsa figure, perception of which also depends on recurrent processing by the grouping of Pacmen inducers. If both recognising the partial object and perceiving the illusory shape depend on recurrent processing, object recognition should vary as a function of the presence of illusory shape. Across two behavioural experiments and a separate EEG decoding study, we found no evidence for an interaction between illusion perception and object recognition, and different neural time courses for the illusory figures alone compared to those of the partial objects, which were decoded earlier. Ultimately, our study highlights the robustness of the visual system to solve the identity of the ambiguous object, independently from the processing of different ambiguities occurring at the same time, providing new insights into the mechanisms of recurrence in the early and late stages of information processing and its application to ambiguous object recognition.

## Introduction

We quickly identify objects in our environment even if they change position, lighting condition, or are seen from a different angle (Biederman, 1995; DiCarlo et al., 2012; DiCarlo & Cox, 2007; Wardle & Baker, 2020). Accurate object recognition occurs in a fraction of a second (Carlson et al., 2013; Hung et al., 2005; Robinson et al., 2023; Thorpe et al., 1996), which has led to proposals of models with purely feedforward processes, consisting of a sequence of feedforward stages arranged hierarchically (Biederman, 1987, 1995; DiCarlo et al., 2012; DiCarlo & Cox, 2007; Riesenhuber & Poggio, 1999; Serre et al., 2007). Although object recognition is highly robust, it becomes more challenging when visual input is ambiguous (e.g., when the object is partially occluded), and yet, we still manage to recognise objects when they are only partially visible. It is understood that object recognition in these circumstances depends on recurrence in the visual system, that is, on lateral and feedback connections that enable information in higher areas to compensate for the ambiguity inherent in the feedforward information (Felleman & Van Essen, 1991; Kietzmann et al., 2019; O’Reilly et al., 2013; Teichmann et al., 2021, 2022). Recurrence occurs across different brain areas along the ventral stream and at different stages of information processing (Cichy et al., 2014; Rajaei et al., 2019; Wyatte et al., 2014). Two processes that evidence this recurrence are grouping and filling-in, and studying them together would further the understanding of how recurrence works when the same information is ambiguous due to different characteristics of the object (e.g., being occluded by an illusory ‘occluder’).

A high-level example of a neural mechanism through which ambiguous object information is resolved is the filling-in process. In this paper, we refer to filling-in as the neural mechanism that reconstructs the missing parts of an object that is partially occluded, which requires feedback connections to the lateral occipital cortex (LOC) (Chen et al., 2020; Ernst et al., 2019; Rajaei et al., 2019; Tang et al., 2018). Another example of neural mechanisms to resolve ambiguous information is grouping. Grouping occurs in an early stage of visual perception, where the brain automatically binds individual elements and perceives them as a whole object, or when closing the contour of a shape (Rock & Palmer, 1990; Roelfsema, 2006; Ullman, 1987). The grouping mechanism in the brain results in the perception of illusory contours, such as in Kanizsa figures, which are a result of arranging Pacman shapes in a way that induces the illusion of a figure without real contours (Kanizsa, 1976). The perception of these illusory contours, in contrast to real edges, occurs due to feedback processes to the primary visual cortex (V1) (Kanizsa, 1976; Lee, 2003; Lee & Mumford, 2003; Lee & Nguyen, 2001; Pak et al., 2020). Previous work shows that grouping and filling-in processes rely on connections to different areas along the ventral stream, such as V1 or LOC, respectively (Lee & Nguyen, 2001; Tang et al., 2018), and affect different stages of processing (O’Reilly et al., 2013), suggesting that they have separate underlying neural mechanisms. However, it is possible that the perception of the occluder, caused by the illusory contours, may facilitate the recognition of the object that is occluded, causing these mechanisms to interact. Following this idea, it would be easier for the visual system to recognise a partially visible object that appears to be behind an occluder, compared to one that is simply missing a portion. However, we do not know what would happen if the occluder itself is also inherently ambiguous enough to require additional recurrent activations, either the same or different, to be perceived. Here, we use both illusory contours and occlusion of objects simultaneously to investigate how grouping and filling-in interact during the processing of partial object images. Our study provides insight into whether the same neural mechanisms underpin both processes, and how these mechanisms would resolve the ambiguous information that arises as a result of grouping and filling-in to recognise the object.

Recurrence supports object recognition by complementing initial feedforward connections by filling in the missing information in previous stages of processing (Kietzmann et al., 2019; O’Reilly et al., 2013; Rajaei et al., 2019). One of the differences between feedforward and recurrent processing is their timing, as recurrence builds upon feedforward processing and takes more time in general (Wyatte et al., 2014). This difference in timing, at the scale of milliseconds, is the reason we need high temporal resolution methods, such as electroencephalography (EEG). More specifically, feedforward processes occur after 40 milliseconds in V1 up to 100 milliseconds in the LOC (Cichy et al., 2014; Rajaei et al., 2019), followed by feedback connections starting with 10 ms of delay in the immediate lower level (Wyatte et al., 2014). Overall, these processes result in recurrent activation in V1 around 80 to 100 milliseconds after the stimulus onset (Boehler et al., 2008; Corthout et al., 1998; Fahrenfort et al., 2007, 2008; Koivisto et al., 2011). For the perception of illusory edges, recurrent activation to V1 happens around 100 ms after the stimulus onset (Lee, 2003; Lee & Mumford, 2003; Lee & Nguyen, 2001), for the illusory shape to be then completed in the LOC (Chen et al., 2020; Murray et al., 2002, 2006). On the other hand, the filling-in process that occurs as a result of occlusion needs recurrent mechanisms to LOC around 130 ms after the presentation of the stimulus (Hegde, 2008; Lerner et al., 2002; Rajaei et al., 2019). The mechanisms used for the completion and filling-in seem to initially differ in time and location, but might overlap at some point of the ventral stream (completion of the filling-in and the completed illusory figure happening in LOC). Consequently, combining EEG with decoding analyses will show what information is in the brain at different points in time, with a resolution of milliseconds (Carlson et al., 2019; Grootswagers et al., 2017; King & Dehaene, 2014). This enables the study of the time courses of object recognition, as well as the underlying mechanisms of grouping and filling-in, when these two mechanisms are required to recognise the same object, in the context of perceiving the illusory shape as an occluder before filling in the occluded object.

In the current study, we investigate the interaction of recurrence in the early and late stages of object recognition through both the use of the perception of illusory edges by grouping mechanisms and the filling-in of occluded information. To examine these mechanisms simultaneously, we present partial images of objects effectively ‘occluded’ by a figure defined by illusory contours caused by aligned inducers in two behavioural experiments (a category and an object task) and one neuroimaging experiment. With this approach, we can gain insights into the underlying processes of grouping and filling-in when they are applied to object recognition at the same time. One possibility is that the presence of the illusory shape ‘occluding’ the object would facilitate its recognition, leading to better behavioural performance and stronger object encoding in the brain (Johnson & Olshausen, 2005; Rauschenberger et al., 2004). Alternatively, if filling-in and grouping share mechanisms, perceiving the ‘occluding’ illusory shape resulting from grouping the inducers (Kalar et al., 2010; Kellman, 2006; Kellman et al., 1998) would compete for recurrent resources, thus decreasing behavioural performance and object encoding of the partial object image. Or, lastly, the feedback processes may not interact (Spehar & Halim, 2016), thus not affecting behavioural performance or encoding strength of the objects. To disambiguate these options, we need to look at the simultaneous presentation of illusion and occlusion in the context of object recognition to understand the interaction of the different types of ambiguous non-trivial object recognition. Our results indicate that illusory contours and the recognition of ‘occluded’ objects do not interact. This suggests independent mechanisms that account for the perception of the illusory contours and the filling-in of the image of the object when it is partially present. Our study sheds light on how recurrent mechanisms in the brain at early and late stages of object processing can work independently, even when they are needed to recognise the same ambiguous object.

## Methods

### Stimuli

To study the neural dynamics of grouping and filling-in, and how these processes impact recognition of partial objects, we began by sourcing 90 greyscale images from pngimg.com. To ensure variability among the types of objects, there were 10 categories (e.g., furniture, tools, insects, mammals) of equal numbers of biological or non-biological objects, each containing three different objects (e.g., glasses, hats, socks). Each object had three different exemplars, for a total of 90 different images (see Figure 1, panel A).

**Figure 1.**
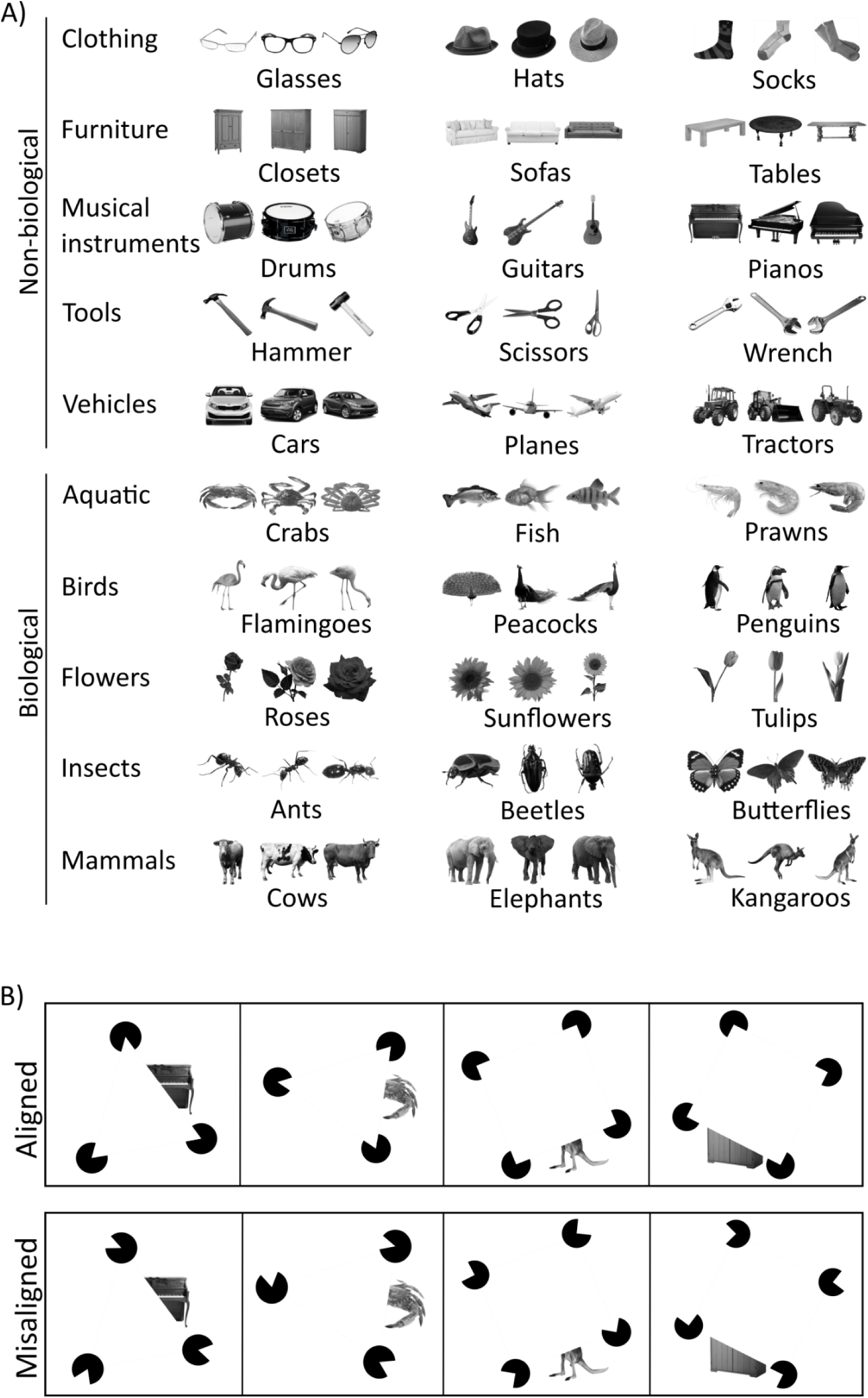
Categories of the objects and conditions used. A. The stimulus set comprised greyscale images of biological and non-biological objects. There were 90 images in total, 3 per object. B. Object images were presented in partial format to create the occlusion of the object. On each trial, the Kanizsa inducers could be aligned or misaligned, resulting respectively in the perception of an illusory triangle/square or not. The position of the inducers in both the misaligned condition and the illusory shapes (triangles and squares) was randomly rotated between images. The inducers and stimuli in this figure are not to scale.

We had two inducer Alignment conditions, which respectively served to either induce the perception of an illusory shape or break it (i.e., valid or invalid illusion). To create the stimulus set, each image appeared in the same partial format in both Alignment conditions (see Figure 1, panel B). To control for effects due to the low-level features from the inducers (e.g., the presence of inducers in the same position and only forming one figure could eventually give away the illusion without the visual system perceiving the illusory contours), the Kanizsa figures comprised either triangles or squares with aligned or misaligned inducers, thus creating or breaking the illusion of contours, respectively. Each unique image was either combined with a triangle or a square in a pseudo-random way, with two images from the same object matched to one shape and one image from the same object matched to the other shape. For each category, there were an equal number of items with triangles and squares. Each Kanizsa figure was randomly rotated, ensuring that only one edge of the object was in line with the inducers, creating the perception of being caused by the occlusion, so the edges of the object would not give away the contours of the Kanizsa figure. The inducers of the illusions were also randomly rotated in the misaligned condition.

In addition to the partial object images, we also included full (non-cropped) images of the objects in the context of the same Alignment conditions described above. This created the perception of the objects being ‘in front’ of the illusory shape. We did this to account for a potential interaction between the position of the object (occluded and in front) and the Alignment condition, if there had been an effect caused by perceiving an illusory shape. However, since we did not find an effect of the Alignment of the inducers, no further analyses with full images were necessary.

### Online behavioural experiments

We conducted two behavioural experiments to assess how well observers can categorise (Experiment 1) and recognise (Experiment 2) full and partial objects presented in the context of aligned and misaligned Kanizsa figures. Both experiments were conducted online (Grootswagers, 2020), hosted on Pavlovia (Peirce et al., 2019), and delivered via a web browser (De Leeuw, 2015). Participants for both experiments were recruited via Western Sydney University’s SONA system, provided informed consent and were compensated with course credits. To study different stages of object recognition, the task of the first experiment was to categorise whether the images were biological or not, while for the second experiment, the task was to recognise each object.

In Experiment 1 (n = 138, where 112 were female and 2 were non-binary, with a mean age of 23.9 years, standard deviation of 8.3), each stimulus was presented for 100 ms, followed by a blank screen until the participant responded. The task was to categorise the stimulus as a biological or a non-biological object using the ‘m’ and ‘z’ keys, which were counterbalanced between participants. Each stimulus was presented once, making it a total of 360 trials.

For Experiment 2 (n = 61, where 37 were female and 2 were non-binary, with a mean age of 30.8, standard deviation of 12.2, a different sample from the previous experiment), the participants saw a word for 500 ms, followed by a stimulus for 100 ms. The participant’s task was to evaluate whether the stimulus represented the same object as the word they had just seen, with half of the trials being the same and the other half different. The different words were always drawn from within the same category as the stimulus (e.g., furniture, tools). This experiment also consisted of 360 trials, one per stimulus.

### EEG experiment

Participants for the EEG experiment were also recruited through Western Sydney University’s SONA system, but did not also participate in the behavioural experiments. One participant was excluded due to a technical error, leaving a final sample of 29 participants for the analysis (23 female and 1 non-binary, mean age of 28.3, standard deviation of 7.8). Participants provided written consent and filled out a demographic questionnaire, where we ensured they had normal or corrected-to-normal vision and did not have a history of photo-sensitive epilepsy. None of the participants were excluded. After the experiment, they received compensation of 6 course credits or a $30 AUD voucher.

EEG Participants sat in a dimly lit, electrically shielded booth at approximately 50 cm from a 24-inch ViewPixx monitor (VIEWPixx /EEG, VPixx Technologies). The participants were instructed to fixate their eyes on a bullseye point (54 x 54 pixels, approximately 1.4° visual angle) in the middle of a mid-grey screen while seeing a fast-rate stream of the stimuli (513 x 513 pixels, approximately 15.4° visual angle, in 2.5 Hz sequences, 200 ms on, 200 ms off). This rapid design has been extensively validated in previous work to evoke reliable neural object representations (Grootswagers et al., 2019, 2022, 2024; King & Wyart, 2021; Mohsenzadeh et al., 2018; Robinson et al., 2019, 2021, 2025). The stimuli that appeared in each sequence consisted of both full and partial objects with aligned and misaligned Kanizsa figures, in a randomised order. Each sequence was composed of 180 stimuli out of the 360, with a total of 24 sequences, which resulted in 12 repetitions of each stimulus for the 4 conditions. Unlike the behavioural experiments, here participants performed an orthogonal task of counting how many Kanizsa figures appeared without an object during each sequence. They responded after the presentation of each stream, using the numbers on the keyboard in front of them. There was a random number of these targets per sequence, varying between 2 and 8. Targets only served to encourage participants to pay attention to the stimuli and were discarded from further analysis of the data. The experiment duration was approximately 40 minutes since the number of targets varied and the participants could take self-paced short breaks between sequences.

### EEG recording and preprocessing

The EEG data were continuously recorded using a 64+2 electrode set with a BioSemi EEG system at a sampling rate of 2048 Hz (BioSemi, Amsterdam, The Netherlands). We followed the International 10-20 system for electrode placement (Jasper, 1958; Oostenveld & Praamstra, 2001), and kept voltage offsets under 20 mV. We pre-processed the signal using the MNE library (Gramfort et al., 2013; Larson et al., 2024) in Python. First, we re-referenced the data to the common average, and we applied a 0.1 Hz high-pass and a 100 Hz low-pass filter. Then we epoched the raw EEG data from -100 to 800 ms, and they were baseline-corrected from -200 to 0 ms before the stimulus onset. Lastly, we down-sampled the data to 1000 Hz. We followed these steps for each participant using all the channels from the EEG signal at every time point. We used the Brain Imaging Data Structure format (Gorgolewski et al., 2016) for EEG data files (Pernet et al., 2019).

### EEG Decoding Analyses

We began by running a verification analysis to determine if the EEG signal contained information about the presence of an illusory shape. To achieve this, we decoded the Alignment of the inducers, training a classifier to distinguish whether the inducers presented onscreen were aligned (thus forming a valid illusory shape) or misaligned (no illusory shape formed), which required generalisation across the different illusory shapes (triangle or square) and rotations of the illusion and the inducers, since we trained and tested across both shapes and all the rotations. This step is important because, after making sure that the neural responses do have information about the presence of illusory contours, we can infer 1) that any effect of Alignment is due to the perception of the illusion, or 2) in the case that we do not find an effect of the Alignment, the absence of perception of an illusion is not the reason for it. Additionally, we can compare the time course of the decoding of the illusion with the main decoding (object recognition and categorisation into biological and non-biological) to consider the order in which the brain processes the information.

For the main EEG analyses, we used decoding to investigate the time course of the neural mechanisms of ambiguous object recognition at the early and late stages of information processing (Grootswagers et al., 2017). To study different parts of the object recognition process, we ran separate decoding analyses at the level of object identity (e.g. kangaroo vs. table vs. shoe, etc), and object category (e.g., biological vs. non-biological) when the image was partial (see Figure 1). For the object identity, we decoded each object presented in the experiment (i.e., 1/30). Using three exemplars per object allowed us to control the differences between the low-level features of each image as much as possible, as the object decoding could now be attributed to the recognition of the object in each image that belonged to that object category. On the other hand, the category decoding entailed the classification of the objects as biological or non-biological (i.e., 1/2), occurring in later stages of the information processing. This allowed us to look into a more abstract representation of the objects, as well as into mechanisms resulting from feedforward and recurrent processes in different time courses and stages of the object processing.

For all decoding analyses, we used a Linear Discriminant Analysis classifier to divide the data using a linear combination of EEG patterns. We used cross-validation to measure the performance of the classifier, after training it on all the sequences but one and testing it on the remaining sequence. We iterated this process 24 times in a way that the classifier was tested on all of the sequences.

### Statistical Analysis

We analysed the behavioural data of the two online experiments separately using Bayesian statistics in R (Morey et al., 2018). We analysed each participant’s accuracy scores and response times of the correct trials on the tasks, excluding any participants whose accuracy rate was below 60% and had a mean reaction time of over 1 second (resulting in a final n = 112 for the category task, and n = 55 for the object task). To know if they performed better when the inducers were aligned or misaligned in each experiment, we used Bayesian paired-sample t-tests to assess the evidence for the null hypothesis (no difference between the aligned and misaligned conditions) and the alternative hypothesis (difference between conditions). We ran the paired-sample t-test to analyse the behavioural accuracy and reaction times when the images were partial in each Alignment condition. Furthermore, to analyse the decoded EEG data, we applied Bayes Factors to the decoding of the conditions against chance, and the difference between the decoding performance when the inducers were aligned and when they were misaligned.

For the Bayesian analyses that test whether the decoding accuracy of each condition was above chance, we used a one-sample t-test with a half-Cauchy prior to make the test directional and used the default width of *r = 0.707*. We centred the null at 0, and we used a prior range from *δ = 0.5 to Infinite* to effectively account for small effect sizes as evidence for the null hypothesis too (Teichmann et al., 2022). To analyse the effect of inducer Alignment over time, for both the decoding and behavioural data, we calculated a paired-sample t-test, pairing the aligned and misaligned inducer conditions. We applied a full-Cauchy prior, as we had multiple possible hypotheses about the direction of the effect. We used the default prior width of *r = 0.707,* and the prior ranges from *δ = -Infinite to -0.5* and *δ = 0.5 to Infinite* for consistency with the other analyses. We used Bayes Factors because of their ability to indicate the strength with which the data supports the alternative (a difference in decoding and participant’s performance when the inducers were aligned, forming an illusory shape, compared to when they were misaligned) or the null hypothesis (no difference between aligned and misaligned inducers).

## Results

### Behavioural experiments

In experiments one and two, we studied how grouping and filling-in processes were translated into behavioural recognition and categorisation of objects, to further compare them with the neural data. We tested whether participants’ ability to explicitly recognise partial objects was modulated by the perception that an illusory shape occluded the objects.

For Experiment 1, neither accuracy nor reaction times appeared to differ systematically between the aligned or misaligned inducer conditions. When participants categorised partial images of objects as biological or non-biological, the Bayes Factors suggested evidence for the null hypothesis for accuracy (BF_10_ = 0.39, error (%) = 8.98 × 10⁻¹¹) and reaction times (BF_10_ = 0.38, error (%) = 2.2 × 10⁻^5^) (see Figure 2. A). These results show no evidence that observers’ ability to categorise the biological status of partial object images varied as a function of the Alignment of the inducers.

**Figure 2.**
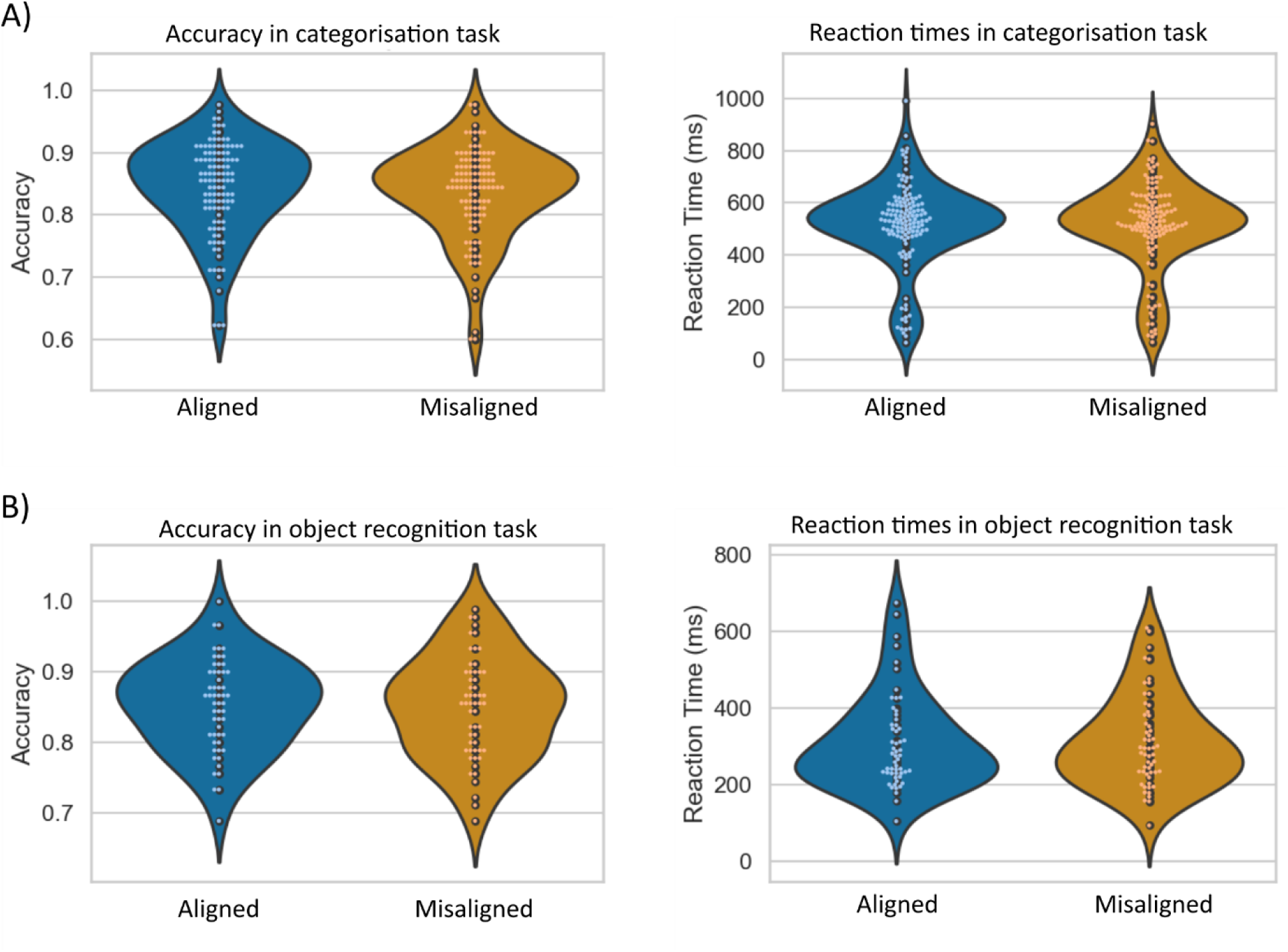
Behavioural results. Accuracy and reaction times are shown as a function of inducer alignment for A. Experiment 1, in which participants judged biological status, and B. Experiment 2, in which participants judged whether an object image matched a preceding label.

Following this, the accuracy and reaction times of the participants in Experiment 2 also showed no difference between aligned and misaligned inducers. As in Experiment 1, a paired t-test between aligned or misaligned inducers showed evidence in favour of the null hypothesis for accuracy (BF_10_ = 0.53, error (%) = 7.33 × 10⁻^7^) and reaction times (BF_10_ = 0.52, error (%) = 1.62 × 10⁻^6^) (see Figure 2. B). In line with the findings from Experiment 1, these results did not show any evidence for the Alignment of the inducers affecting the participants’ performance in recognising the objects.

### EEG experiment

Before examining how information about objects in the EEG signal varied depending on the Alignment of the inducers, we verified that the neural signal contained information about the presence of the illusion itself, formed by the aligned inducers (see Figure 3). The presence/absence of illusory shapes was consistently decodable from 150 ms after the stimulus onset. In general, this suggests that the participants perceived the illusion, which allowed us to look at the effect of the Alignment of the inducers on the occlusion of the images by an illusory shape.

**Figure 3.**
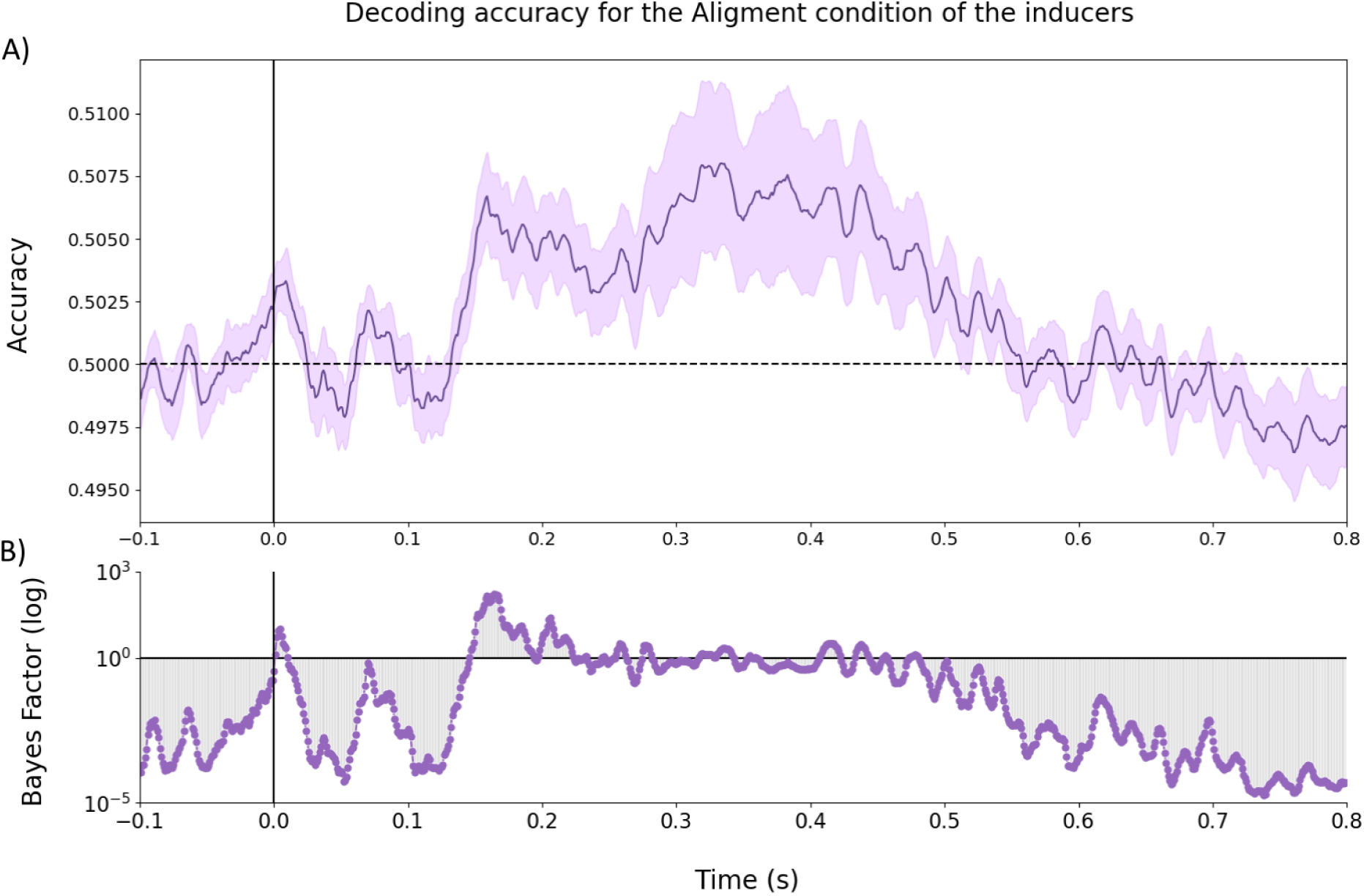
EEG Decoding of Alignment condition. A. Decoding accuracy over time of the perception of the illusion (caused by aligned vs. misaligned inducers). Shaded areas are standard error, and 0.5 represents the chance level. B. Bayes Factors reflecting decoding above 50% chance level across time.

Our main question of interest was to investigate how object representations evoked by partial object images differed depending on whether they were perceived as ‘occluded’ behind an illusory shape caused by the Alignment of its inducers. To this end, we ran a separate decoding analysis at two levels of categorisation for our stimuli (i.e., category = biological vs. non-biological, and object = glasses vs. hats vs. dogs….), contrasting decoding accuracies obtained when the object appeared in the context of a valid or invalid illusion (see Figure 1). The decoding accuracy scores for both object and category decoding (Figure 4. A) for images presented with aligned and misaligned inducers were above chance from approximately 100 ms after stimulus onset. Category decoding of stimuli with aligned inducers remained above chance and stayed more or less constant, whereas accuracy dropped around 350 ms when the image had misaligned inducers (Figure 4. A, left). To quantify the effect of the presence of the illusion caused by the Alignment of the inducers, we calculated the Bayes Factors for the difference in decoding accuracies between the aligned and misaligned conditions (Figure 4. C, left). In this case, there was only evidence for the null hypothesis (i.e., no difference between the aligned and misaligned inducers), with very small and unsustained effects at around 200 and 500-600 ms that were not enough to make a case for a difference. The object decoding followed a similar pattern of results when the stimuli had aligned and misaligned inducers, both staying constant above chance and dropping before 400 ms (Figure 4. A, right). As per the Bayes Factors of the difference between the conditions, there was evidence for a difference in alignment of the inducers around 100-110 ms and after 200 ms (Figure 4. C, right), with higher decoding accuracy for misaligned compared to aligned. However, this effect was also not sustained enough to make a confident case for a difference.

**Figure 4.**
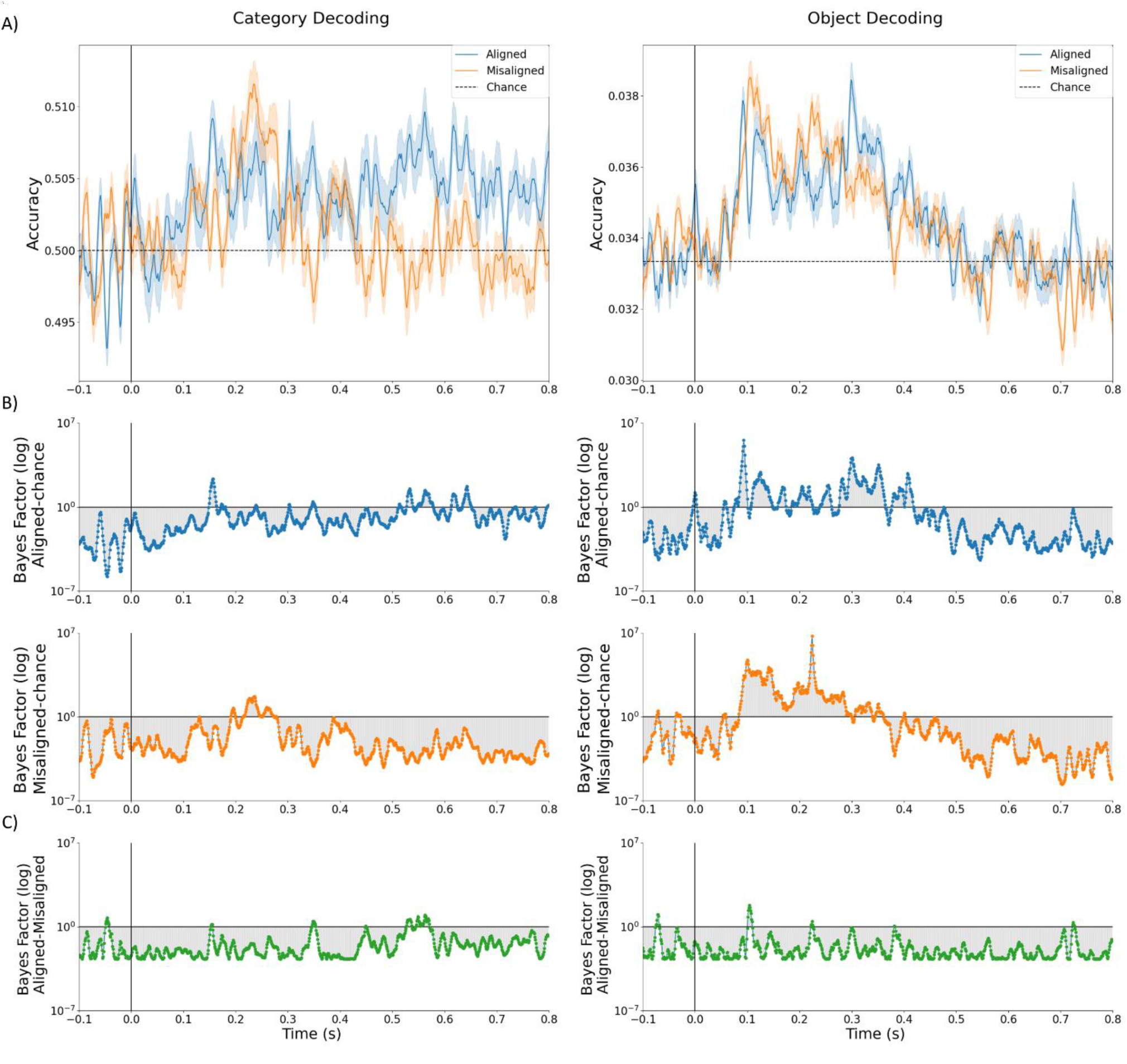
EEG Decoding of categories and objects. A. Category (left) and object (right) decoding accuracy of the classifier when the image was presented behind the illusion over time, depending on the Alignment of its inducers. The blue line corresponds to aligned inducers, whereas the orange line depicts the decoding scores of the misaligned condition. The shaded areas around the accuracy scores are the standard errors from the participants. B. Bayes Factors of either aligned or misaligned conditions against chance. C. In green, the Bayes Factors show how the effect of the difference in Alignment was over time.

## Discussion

This study aimed to assess the early and late stages of ambiguous object recognition, through the mechanisms of grouping and filling-in, by using illusory contours and occlusion of objects, respectively. We studied how the perception of a partial object could be affected by the occlusion with an illusory shape, by manipulating the alignment of the inducers to either form a valid Kanizsa illusory figure or not. We tested participants’ behavioural ability to either recognise the identity or categorise the biological status of partial object images that either appeared to be occluded by an illusory shape or not (as induced by varying the alignment of inducers). For both the categorisation and object identification experiments, we found no evidence that either accuracy or reaction times differed as a function of the illusion’s presence. To examine the neural mechanisms underlying the behavioural results, we collected time-series EEG while participants were looking at a fast stream of stimuli. After decoding the presence of the Kanizsa shapes to ensure that the neural signal contained information about the illusion, we examined object and category representations evoked by the partial object images when presented with aligned or misaligned inducers.

Our decoding results showed a relatively late encoding of the illusory contours (approximately 150 – 250 ms after stimulus onset). In contrast, information about object identity and object category arose earlier in the neural signal than information about illusion presence (after 100 ms onwards). Moreover, these object and category representations appeared to be independent of the illusion presence, as evidenced by the lack of influence that the presence or absence of an illusory shape had on the recognition of the object, both behaviourally and in the neural signal. Decoding the alignment of the inducers demonstrated that the experiment design and stimuli were effective in evoking the illusion when the inducers were aligned, which strengthened our results despite not finding a difference in alignment applied to the recognition of the partial objects.

Our research builds on prior work on the time course of ambiguous object recognition and the recurrent activation of both grouping (Rock & Palmer, 1990; Roelfsema, 2006; Ullman, 1987) and filling-in processes (Ernst et al., 2019; Rajaei et al., 2019; Tang et al., 2018). With the time course of the first grouping’s feedback connections to V1 being 100 ms after stimulus onset (Lee, 2003; Lee & Mumford, 2003; Lee & Nguyen, 2001) and with filling-in happening later, around 130 ms after the stimulus onset (Lerner et al., 2002), we expected the partial object to be recognised after the completion of the illusion, that would serve as an ‘occluder’. Based on previous research on occlusion, the recognition of the object would have resulted in either an interaction caused by a facilitation to fill in the information of the ‘occluded’ object due to the presence of an ‘occluder’, the illusion in our case, (Johnson & Olshausen, 2005; Rauschenberger et al., 2004) or in the competition resulting from the use of the same neural mechanisms to assign the modal and amodal objects (Kalar et al., 2010; Kellman, 2006). In our experiment, although we could decode both the representation of the objects for the partial images and the presence of the illusion that emerged from aligning the inducers, we found that the grouping mechanisms to solve the illusory contours and the recognition of the occluded object might have worked independently to solve the ambiguous information. That is, the grouping that occurred to create the illusion to occlude the objects did not affect the recognition of these objects, as they were perceived earlier than their occluder. We believe one of the reasons this happened was because of separate mechanisms used for the perception of the illusion and the recognition of the objects independently, aligning with studies identifying different neural mechanisms for grouping and filling-in (Spehar & Halim, 2016).

Importantly, the lack of an interaction between illusion perception (due to grouping of the inducers) and object recognition (due to filling-in of the partial object image) does not appear to be a consequence of observers not perceiving the illusory shape at all. Indeed, decoding results for the alignment of the inducers confirmed that there was a difference in the processing of the presence and absence of a Kanizsa shape in our stimuli. Furthermore, the design of the experiment and the stimuli (e.g., fast and randomised presentations, rotations of all the shapes and the inducers, use of squares and triangles) prevented the perception of the illusion from being a consequence of any effect other than the alignment of the inducers. Combining all these measures with above-chance decoding of the alignment of the inducers, and the identification and categorisation of the objects, suggests that our effects are a consequence of mechanisms that might have worked independently, to perceive the illusion and to recognise the objects, respectively. On one hand, the perception of the illusion occurred later than the first 80 ms of visual processing, with an onset of around 150 ms. This is consistent with the assertion that perception of illusory contours requires feedback from extrastriate areas to earlier visual areas, and that its decoding onset reflects the encoding of the completed illusory shape in LOC (Chen et al., 2020; Hegde, 2008; Murray et al., 2002, 2006; Lerner et al., 2002). On the other hand, however, when we examined the object representations in the context of the grouping mechanism that supports the perception of illusory contour, the information of the object in the brain had an onset of around 100 ms, approximately 50 ms before the processing of the illusory ‘occluder’. Although both object recognition and its categorisation peaked after the recognition of the ‘occluder’, we found no evidence of a difference between aligned and misaligned inducers while recognising and categorising the object for both decoding and behaviourally. Thus, supporting the independence of the perception of the illusory figures and the recognition of the object.

The representation of the objects having an earlier onset time compared to the perception of the illusory shapes as occluders might have been caused by the stimuli we used in our experiment (combining familiar objects with Kanizsa figures). That is, the objects used here might have remained very recognisable despite the presence of the illusion and their partial presentation, which produced high performance in behaviour responses and decoding accuracy, possibly as a result of no filling-in being necessary. This could be the reason for the lack of influence that the presence of the illusion had on the objects, since the illusions were more ambiguous than the recognition of the objects on their own. Following our results, we believe the recognition of the partial object image might have occurred throughout the initial processing of the object, independently of the resolution of the illusory shape as an occluder, and possibly without a filling-in process. A possible explanation for this is the difference in familiarity between the objects and illusions, creating a difference in ambiguity between the partial objects and the illusory contours, explaining the facilitation of the recognition of the objects compared to the perception of the illusions (Hazenberg et al., 2014; Hazenberg & van Lier, 2016; Helton & Nanay, 2019; Vrins et al., 2009; Yun et al., 2018). Future studies could solve the difference in ambiguity by ensuring that the grouping effect happens before the object recognition, either by presenting the illusion 100 ms before the occlusion takes place or increasing the ambiguity of the objects by using different types of stimuli (e.g., line drawings, Gabors).

Alternatively, our results might be due to the perception of depth in our stimuli not being correctly assigned by the visual system, as the illusory shape was supposed to be perceived in front of the object. Thus, the visual system would not be able to solve the extrinsic contours of the ‘occluder’ before the recognition of the object (Anderson, 2007; Kogo et al., 2014; Kogo & Wagemans, 2013; Nakayama et al., 1989), resulting in not being able to designate the illusory shape as ‘occluder’, as we expected. This lack of pairing in the brain between the illusion and the occluder might have resulted in independent processes to recognise the objects and to perceive the illusion, as our findings indicated. It would be interesting to further analyse the depth cues caused by the Kanizsa shapes occluding objects.

In conclusion, our research suggests that the mechanisms used to perceive the illusory shapes and to recognise the partial objects operate independently, at lower and higher levels of object recognition, respectively. The data here suggest different neural dynamics underlie the processes to support the recognition of an object ‘occluded’ by a Kanizsa. This has implications for understanding how the mechanisms to perceive the illusion and to recognise the ‘occluded’ objects operate. Ultimately, our study highlights the robustness of the visual system to solve the identity of the object, independently from the processing of different ambiguities occurring at the same time, such as illusory contours. This has significance to gain insights into the interaction of different mechanisms in the early and late stages of information processing and their application to ambiguous object recognition in other areas, like machine vision.

## Data and code availability

Stimuli and analysis code are available here: https://osf.io/x6fv9/. Data will be made publicly available upon publication.

## Acknowledgements

This research was supported by ARC DE230100380 (TG) and ARC DP220103047 (MV). ARH is supported by a MARCS Institute PhD Scholarship.

